# The Neural Underpinnings of Aphantasia: A Case Study of Identical Twins

**DOI:** 10.1101/2024.09.23.614521

**Authors:** Emma Megla, Deepasri Prasad, Wilma A. Bainbridge

**Author notes:** Correspondence to: Emma Megla, Institute for Mind and Biology 940 E. 57th St. Chicago, IL 60637. These authors contributed equally (Emma Megla, Deepasri Prasad).

## Abstract

Aphantasia is a condition characterized by reduced voluntary mental imagery. As this lack of mental imagery disrupts visual memory, understanding the nature of this condition can provide important insight into memory, perception, and imagery. Here, we leveraged the power of case studies to better characterize this condition by running a pair of identical twins, one with aphantasia and one without, through mental imagery tasks in an fMRI scanner. We identified objective, neural measures of aphantasia, finding less visual information in their memories which may be due to lower connectivity between frontoparietal and occipitotemporal lobes of the brain. However, despite this difference, we surprisingly found more visual information in the aphantasic twin’s memory than anticipated, suggesting that aphantasia is a spectrum rather than a discrete condition.

## Introduction

What does your bedroom look like? For many of us, we can form a vivid mental image of this place, filling our “mind’s eye” with its visual details. However, the nature of these mental representations—and their relationship to perception—is debated in the field. Are these representations a recapitulation of what we viewed during perception, or have they been altered in memory? The key to this fundamental question may be those with *aphantasia*, a condition characterized by the lack of voluntary mental imagery (Keogh & Pearson, 2018; Zeman et al., 2015). Since aphantasia may serve as a natural “knock-out” model of visual imagery and recall, it could highlight potential differences in our perceptual and mnemonic representations. Here, we leveraged the identical genetics and shared experiences of a unique case study: a pair of identical twins—one with aphantasia and one with normal imagery—using neuroimaging to pinpoint differences in their memories stemming from their different imagery experiences.

It is currently debated how visual perception relates to visual long-term memory. On one hand, a collection of research has identified similarities between perception and memory, finding that the same voxels activated during perception are reactivated during memory (Cichy et al., 2012; Ishai et al., 2000; O’Craven & Kanwisher, 2000; Reddy et al., 2010). On the other hand, more recent studies have identified meaningful differences between perception and memory, uncovering that the voxels activated during memory are anterior to—rather than the same as— those activated during perception (Bainbridge, Hall, et al., 2021). Moreover, entirely different networks may even be involved in perception than in memory (Baldassano et al., 2016; Silson et al., 2019; Steel et al., 2021). In fact, the existence of aphantasia suggests differences between perception and memory: if memory is a reinstatement of perception, then how do aphantasics have intact perception, but disrupted memory (Dijkstra et al., 2019)?

Although aphantasia could help define the relationship between perception and memory, we first need to understand the nature of this condition. As aphantasia has largely been identified through the Vividness of Visual Imagery Questionnaire (VVIQ; Marks, 1973; Zeman et al., 2020), resulting in estimates that roughly 4% of the population has aphantasia (Dance et al., 2022; Zeman et al., 2015), this subjective measure has led to claims that it could instead be a metacognitive or psychogenic condition (de Vito & Bartolomeo, 2016). Indeed, there is little objective evidence of aphantasia, although measures have been quantified in recent years. For aphantasics, forming a mental image does not bias perception during subsequent binocular rivalry (Keogh & Pearson, 2018), unlike for those with normal imagery (Pearson et al., 2008, 2015). Additionally, those with aphantasia have a reduced skin-conductance response when reading a frightening story compared to controls (Wicken et al., 2021), as they cannot “see” these events in their minds. Therefore, is lack of imagery a *subjective* or *objective* experience? Neuroimaging could help reveal the underlying nature of this condition, and potentially identify additional objective measures.

To date, there have been few published neuroimaging studies of aphantasia, with a majority only published in the last year, and these studies largely taking a network approach (e.g., Cabbai et al., 2024; Chang et al., 2025; Liu et al., 2025; Milton et al., 2021; Weber et al., 2024; Zeman et al., 2010). These neuroimaging studies suggest that aphantasics may have reduced connectivity between their visual-occipital regions and other regions of the brain, such as the prefrontal cortex (Milton et al., 2021) or temporal lobe regions (Zeman et al., 2010; but see Monzel et al., 2024), but increased connectivity among non-visual areas (Milton et al., 2021; Zeman et al., 2010). Similarly, a recent electroencephalography (EEG) study found that mental imagery may be evoked starting in the left temporal lobe for aphantasics compared to frontal areas in normal imagers (Furman et al., 2022). Therefore, it seems that aphantasics may have different networks dedicated to memory than their control counterparts. However, better understanding aphantasia and memory’s relationship to perception will also depend on understanding the neural *representations* during memory. Recent work has started to tackle this topic, though focused on lower-level visual representations (Cabbai et al., 2024; Chang et al., 2025; Liu et al., 2025; Weber et al., 2024).

In the present study, we use functional magnetic resonance imaging (fMRI) to identify neural underpinnings of aphantasia, and examine the nature of memory, through a pair of identical twins—one with aphantasia and one with normal imagery. The identical genetics and shared experiences of these twins ensures that meaningful differences in memory are likely due to their differing imagery experiences, making this an ideal sample to pinpoint neural markers of aphantasia. By having the twins view and mentally imagine the same items, we found that although the aphantasic twin does have lower memory quality, her memories still contained an unexpected amount of visual information. Additionally, we also observed reduced connectivity between occipitotemporal and fronto-parietal areas in the aphantasic twin, likely due to less visual representations even during mind-wandering. Further, we found a marker for atypical lateralization—bilateral language laterality—in the aphantasic twin, which could serve as a mechanism for her aphantasia. These results not only identify some of the first objective, neural measures of aphantasia, but also suggest that memory is more than a recapitulation of perception.

## Methods

### Participants

Two identical twins (31 years old, female) raised in the same household participated in this experiment. Their overall mental imagery abilities were assessed using the Vividness of Visual Imagery Questionnaire (VVIQ; Marks, 1973) and separable object and spatial imagery abilities using the Object-Spatial Imagery Questionnaire (OSIQ; Blajenkova et al., 2006). The twins also self-report as “mirror twins”, in which features such as birthmarks present on opposite sides of each individual. Relatedly, they report opposite handedness (i.e., left-handedness in the aphantasic twin but right-handedness in the imager twin), and so handedness was empirically assessed using the Edinburgh Handedness Inventory (Veale, 2014). The participants had corrected vision and wore MRI-compatible lenses during the scan. Subjects consented to participation, following the guidelines approved by the University of Chicago Institutional Review Board (IRB20-0233), and were compensated for their time.

Additionally, the twins were separately interviewed to gauge their natural history, such as their imagery experiences and interests. Similar to others with aphantasia (Zeman et al., 2015), the aphantasic twin discovered her lack of imagery in her late 20’s when she came across an article online about the condition. Interestingly, the aphantasic twin mentioned that her imagery in other senses (e.g., auditory imagery) is intact, and that she even uses these other senses to compensate for her lack of visual imagery. For example, she mentioned that instead of visually imagining the moves she would make when planning a rock-climbing route, she instead imagines how her weight and body would shift with each move.

The aphantasic twin also shared experiences and hobbies with the imager, such as having the same relationship status at the time of the study (i.e., long-term partner), overlap in their health histories, and some shared hobbies. However, there were also striking differences between them; most notably, the imager twin works in a very visual profession—graphic design after receiving a degree in fine arts—whereas the aphantasic twin works as a chemist. In fact, the imager twin mentioned their attraction towards visually stimulating hobbies, such as animated television and movies, drawing, and calligraphy, whereas the aphantasic twin opts for sci-fi and fantasy media, rock-climbing, and gardening. Therefore, it is possible that the aphantasic twin’s lack of visual imagery affected her life choices and interests, opting for less visually-oriented experiences. All interview questions and the twins’ abbreviated answers are reported in *Table S1*.

### Experimental paradigm

While in the fMRI scanner, the twins primarily performed interleaved runs (4 runs each) of two mental imagery tasks (**Fig. 1**). All tasks performed in the scanner were displayed using Psychtoolbox (Brainard, 1997). The *Novel Imagery* task (**Fig. 1a**) was adapted from Bainbridge, Hall, et al. (2021). In each trial of this task, participants were presented with an image to view for 6 sec. After a 1 sec fixation, they completed a 4 sec distractor where they viewed a series of scrambled images with one intact image and responded when they saw the intact image. After a 1-4 sec jittered fixation, they were instructed to recall the previously shown image for 6 sec as vividly as possible. At the end of each trial, they rated the vividness of their memory as either high vividness, low vividness, or no memory. Participants viewed 48 images of scenes and 48 images of objects presented against a white background, for a total of 96 images. Each run consisted of 24 trials equally divided between scenes and objects. The order of image presentation was randomized, but the same between participants.

**Figure 1.**
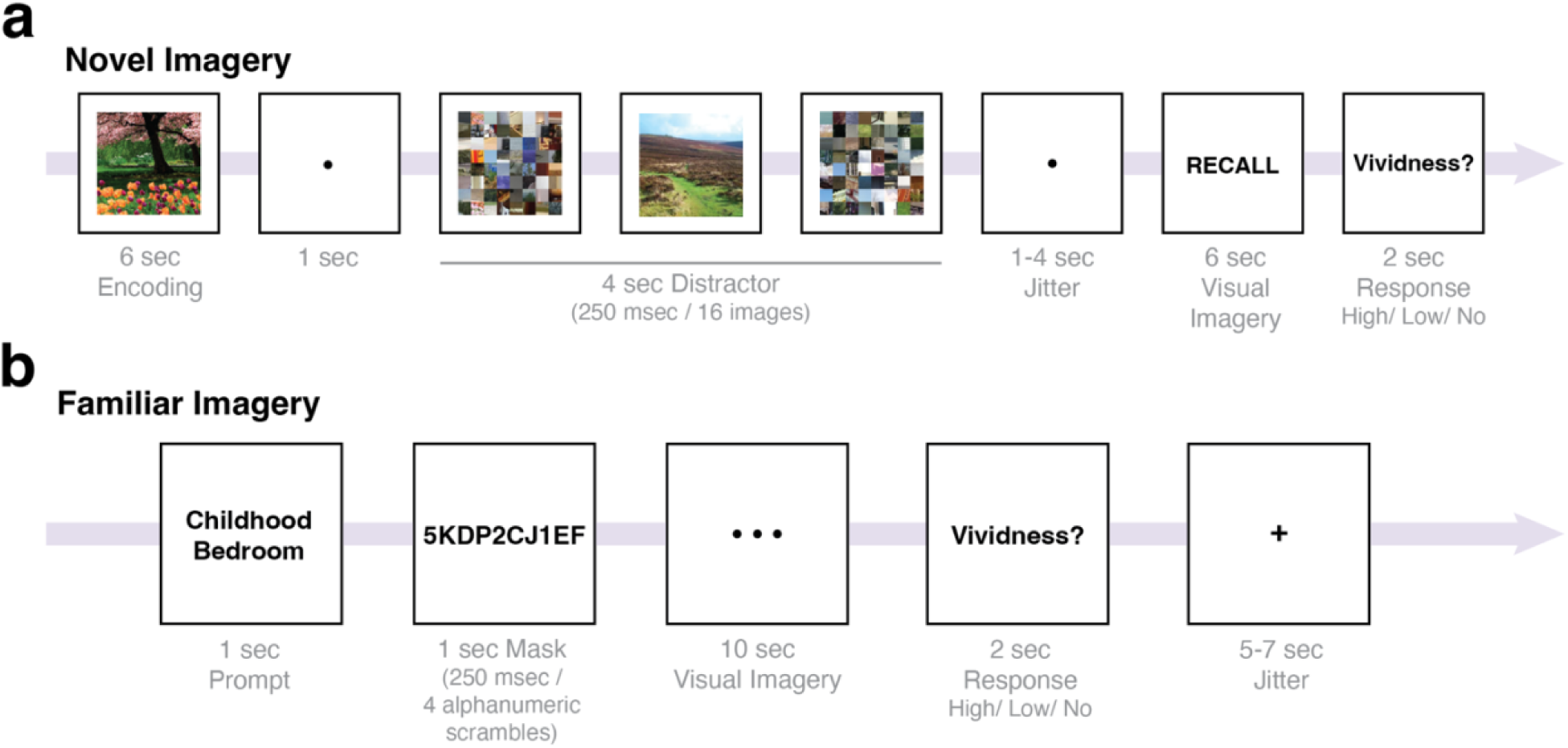
Methods for mental imagery tasks. (a) Methods for the Novel Imagery task. Participants first encoded a novel scene or object image for 6 sec. Then, there was a 4 sec distractor period in which the participants indicated an intact image amongst a stream of scrambled images. After a 1-4 sec randomized jitter, participants then recalled the original image using mental imagery for 6 sec. Lastly, they rated the vividness of their mental image using a three-point scale. There was a total of 96 trials. (b) Methods for the Familiar Imagery task. Participants were first given a prompt which consisted of the text label of a personally familiar person or place. After a 1 sec mask of scrambled alphanumeric characters, the participants then mentally imagined the corresponding text prompt for 10 sec before rating the vividness of their mental imagery using a three-point scale. There was a 5-7 sec randomized jittered fixation between trials and 144 trials total.

The *Familiar Imagery* task (**Fig. 1b**) was modified from Steel et al. (2021). Before the scan, the twins generated a list of 36 personally familiar places and 36 personally familiar people together. In each trial of the experiment, participants were prompted with the name of one of the people (e.g., mom) or places (e.g., childhood bedroom) for 1 sec. After a 1 sec dynamic alphanumeric mask, they were asked to recall the person or place associated with the prompt as vividly as possible for 10 sec. After recall, they were asked to rate the vividness of their imagery as either high vividness, low vividness, or no memory. The trial order was randomized, but the same between the twins. There were 18 trials in each run.

In addition to the two mental imagery tasks, the twins also performed two localizers and a video watching task while in the scanner. Participants completed one run of a perceptual localizer at the beginning of the experiment to identify scene-, face-, and object-selective regions, and involved participants viewing four 16 sec blocks of images. Each block contained a single category of images: faces, objects, scenes, or scrambled images. Participants indicated consecutive repeats of images by pressing the response button.

Halfway through the experiment, they performed one run of a video watching task, which involved watching a 10-min abstract video devoid of semantic content titled *Inscapes* (Vanderwal et al., 2015), which is designed for functional network localization. At the end of the experiment, participants completed one run of a language localizer adapted from (Fedorenko et al., 2010). Given that the twins have different handedness (termed “mirror twins”), it is possible that the twins have different lateralization and organization of their brains. Therefore, by looking into the highly lateralized process of language processing, we can assess whether their whole-brain laterality may differ which could be a potential mechanism behind their differing imagery experiences. In each trial, participants were presented with a 12-unit sequence of either words that formed a sentence or nonwords. Each unit was presented individually for 450 msec. Participants completed 1 run of this task, with 16 blocks of 3 trials of either words or nonwords.

Outside of the scanner, the twins also completed a drawing experiment to obtain visual representations of their perceptual and mnemonic content. During the experiment, participants first encoded three scene images sequentially (a bedroom, living room, and kitchen) for 10 sec each before recalling them using drawing. The canvas used to create the drawings matched the size of the encoded images (500 × 500 pixels). While drawing, participants had access to a range of colors, an eraser tool, and an undo tool. Participants next performed a short old/new recognition task with the three target images and three foil images from the same scene categories. Lastly, the participants were sequentially shown the original three scene images alongside the drawing canvas. The participants were instructed to copy each image using drawing.

### MRI acquisition

Neuroimaging data was collected at the University of Chicago using a 3T Philips Achieva MRI scanner with a 32-channel phased-array head coil. Anatomical scans used a T1 MPRAGE structural scan with a resolution of 1×1×1 mm voxels. Functional scans used a gradient echo-planar T2* sequence (39 axial slices parallel to the anterior commissure-posterior commissure line; 64×64 matrix; FoV=192×192 mm; TR=2000 msec; TE=28; 0.5 mm gap; flip angle=77 degrees; 3×3×3 mm voxels). We preprocessed the functional scans using the same protocol as prior studies (Bainbridge, Hall, et al., 2021), which included slice time correction and motion correction using the Analysis of Functional NeuroImages (AFNI) software (Cox, 1996). No spatial smoothing was applied. Functional data were aligned to Montreal Neurological Institute (MNI) space.

### MRI analysis

#### Whole-brain univariate analyses

We ran general linear models (GLM) to perform whole-brain univariate analyses on the Novel Imagery and Familiar Imagery tasks. For Novel Imagery, all trials were modeled separately (e.g., recalling farmhouse #4). Whole-brain *t*-contrasts were then calculated by grouping trials along the dimensions of object/scene and encoding/recall. Distractor periods were modeled separately. For Familiar Imagery, all trials were also modeled separately (e.g., Student Art Gallery), with trials grouped along the dimensions of recalling people/places for whole-brain *t*-contrasts. Both GLMs additionally included six regressors for head motion. The individual trial beta values were used for multivariate analyses for both imagery tasks.

#### Defining regions of interest

From the Familiar Imagery task, we identified “familiar memory regions” that have been found in the medial parietal area using a people>places contrast. We compared the coordinates of these regions in MNI space to where they have been localized previously (Bainbridge & Baker, 2022; Silson et al., 2019).

#### Identifying peak voxels in univariate activity

To determine whether there was an anterior shift from perception to memory in category-selective areas, we used data from the Novel Imagery task as the twins perceived and remembered the exact same scenes and objects. Therefore, differences in the location of peak voxel activity of these regions from perception to memory would not be confounded with different image sets, and they could instead reflect differences in content when representing these images (e.g., memory semanticization).

When determining whether an anterior shift had occurred between perception and memory in either twin, we focused analyses on perceptual and memory variants of the scene-selective area parahippocampal place area (PPA). The location of peak voxel activity within PPA was determined in MNI space using a scenes>objects contrast to identify the functional region within the posterior parahippocampal cortex (Pruessner et al., 2002). We first used a threshold of p<0.001, but iteratively lowered the threshold until the area was identified. We determined peak voxel activity separately for perception and memory. Note that although PPA is classically defined as a region sensitive to scene *perception*, we are also using that term for any nearby regions sensitive to scenes during recall and imagery. Although PPA was largely identified using the most conservative threshold, the aphantasic’s left and right PPAs were notably not identified during memory until we used a more liberal threshold (left: p<0.02; right: p<0.01). However, we also found similar evidence for an anterior shift when using a more conservative threshold and expanding to the greater medial temporal lobe region (see *Fig. S3*). The same steps were performed using a scenes>objects contrast to identify peak voxel activity in occipital place area (OPA) and medial place area (MPA), as well as an objects>scenes contrast to identify peak voxel activity in object-selective area lateral occipital (LO; see *Table S3* for coordinates of peak voxel activity in these additional regions).

#### Whole-brain SVM searchlight analyses

We performed four whole-brain SVM searchlight analyses to determine representational differences between participants and tasks during the Novel Imagery task. Between-participants, we used the voxels within each searchlight (sphere radius=3 voxels) to train an SVM to differentiate between objects and scenes in the imager and tested the model to differentiate objects and scenes in the aphantasic. We did this twice: first for their representations during perception, and second for their representations during memory. If there are similar representations between the imager and aphantasic in a searchlight area, then there will be higher (and above chance) decoding accuracy. Significance at each searchlight area was determined through permutation testing, in which we performed 100 iterations of randomly swapping half of the scene and object labels during training, and swapping those same labels during test to build a null distribution. We used the same logic within-participants (but between tasks), where we trained an SVM on a participant’s perception and tested on her recall. We performed this for both the aphantasic twin and the imager twin.

We additionally performed a second set of permutation tests to determine searchlight areas that had significantly different decoding accuracy for either (1) encoding or recall or (2) for the imager or the aphantasic twin. We ran this permutation test by randomly swapping half of the condition labels (e.g., encoding or recall) between training and test. We performed this random swap 100 times to build a null distribution, with significance set at p<0.05 for all permutation tests. For all SVM searchlight results, statistics and brain visualizations are shown using the uncorrected threshold of p<0.001, but we find the same trends and key brain regions when using cluster threshold correction (*see Supplemental Results 1 and Fig. S1*). For cluster threshold correction, we performed 1-sided thresholding for the initial SVM searchlight analyses and bi-sided thresholding when comparing between the SVM searchlight conditions (i.e., when results could be positive and negative). Significant areas of interest from this analysis were identified using well-known anatomical landmarks, such as the PHC being on the posterior half of the parahippocampal gyrus (Pruessner et al., 2002).

#### Representational similarity analyses and discrimination indices

We conducted representational similarity analyses (Kriegeskorte et al., 2008) on the Familiar Imagery task data to determine the amount of visual information present in memory when recalling familiar people and places. We performed these analyses in the PHC region identified from the whole-brain searchlight SVM analyses as having unexpectedly higher similarity between perception and memory for the aphantasic than the imager twin. We built a representational similarity matrix (RSM) for each twin by correlating (Pearson’s correlation) the activation of each voxel of the PHC region between each pair of trials. Therefore, trials with a higher Pearson’s correlation have more similar representations.

If there is visual information present during memory of familiar people and places, then we would expect more similarity for within-category trials (e.g., within-people) than between-category trials (i.e., between people and places) as there would be more shared visual information within-categories. Therefore, we calculated discrimination indices (*D*) by subtracting the average of all between-category correlations from the average of all within-category correlations (Kravitz et al., 2011). To determine whether there was any significant difference in the discriminability of people vs. places between participants, we performed a permutation test by calculating the difference in *D* between participants when half of the trials were randomly swapped between participants. We performed 1000 iterations of this random swapping to build a null distribution to compare to the true difference in discriminability between participants.

#### Functional connectivity analysis

To determine participants’ functional connections in the brain at rest, we had the twins watch a 10-min video titled *Inscapes* (Vanderwal et al., 2015) that contained no semantic or social information while in the scanner. The functional data were preprocessed and analyzed using a separate pipeline from the other collected MRI data to more closely follow recent studies of functional connectivity (Chamberlain et al., 2024; Zhang & Rosenberg, 2024). Preprocessing involved using *afni_proc* to remove outliers, perform time-slice correction, align to the anatomical scan, register volumes to the TR with the least amount of motion, and align to MNI space. Additionally, white matter and CSF masks were created for each participant using FMRIB Software Library (FSL) and were regressed out of the data. To control for motion, volumes in which 5% of the voxels contained motion outliers were removed as well as volumes following a change of at least 0.2 mm of motion from the volume. As this last step left too few volumes for analysis for the aphantasic twin, we used a slightly more liberal threshold of censoring out volumes with at least 0.3 mm of motion for the aphantasic twin.

For the functional connectivity analysis, we parcellated the pre-processed data into the 268 nodes of the Shen Brain Atlas (Shen et al., 2013). Since analysis depended on directly comparing the strength of functional connections between the twins, we removed three nodes (2 prefrontal nodes, 1 temporal node) that were filtered out during preprocessing in at least one participant. After calculating the mean time series for each node, we used Pearson’s correlation to correlate the mean times series between each pair of nodes and create a functional connectivity matrix. Lastly, to compare the strength of connections between participants on the lobe level, we averaged across the correlation values within each lobe (i.e., averaged across the nodes) and subtracted these averages between participants (imager twin – aphantasic twin).

#### Mind-wandering model

We used a model created by Ke et al. (in prep) which predicts the content of mind-wandering. This model was created by having typical individuals indicate the content of their thoughts during rest. Specifically, amongst other questions, they reported whether their mind-wandering was in the form of images. The authors built models to predict participants’ self-reported imagery ratings from their functional connectivity data. The model used here included 705 connections that reliably predicted more imagery and 507 connections that reliably predicted less imagery in their data. For each twin, we calculated the strength of this imagery network by subtracting the average strength of the 507 connections predicting less imagery from the average strength of the 705 connections predicting more imagery. Statistical significance was assessed by comparing the actual difference in network strength between the twins to a null distribution of difference values generated by shuffling the edge positions in the imagery network 1000 times before computing network strength. Note that the paper for this model also reports on the twins’ data reported in the present paper.

#### Language localizer analysis

We ran a language localizer analysis to test whether the twins’ language areas were lateralized to different hemispheres, which could be a marker for different brain-wide lateralization and serve as a mechanism for their differing imagery experiences. We determined language lateralization using the previously established method of calculating a laterality index (LI; Binder et al., 1996; Desmond et al., 1995) which involved comparing the number of voxels active in the right (RH) versus the left hemisphere (LH) for words>nonwords at a threshold of p≤0.001. Specifically, we followed the formula LI=(LH–RH)/(LH+RH) and did not include voxels within the cerebellum, as the cerebellum can show opposite trends to the rest of the brain (Hubrich-Ungureanu et al., 2002). Therefore, a positive LI indicates laterality towards the left hemisphere, whereas a negative LI indicates laterality towards the right hemisphere. However, we set a laterality threshold of 0.2 following the most common convention (Deblaere et al., 2004; Springer et al., 1999), which meant that laterality was considered bilateral until the LI was > +/- 0.2.

#### Mental imagery ROI SVM analysis

To determine whether the SVM searchlight results replicate using a region of interest (ROI)-based approach, we used the tool *Neurosynth* (Yarkoni et al., 2011) to localize a mental imagery ROI. When given a term, Neurosynth performs a meta-analysis on all published fMRI studies in its database that includes that term in the abstract, and then determines the voxels that are preferentially active for that term. These voxel maps are then corrected using a false discovery rate (FDR) of 0.01. Therefore, this is a powerful method of deriving ROIs based on data from many studies.

For the present ROI, we used the term “mental imagery”, which created a brain map based on 84 published studies. We focused on replicating the SVM searchlight results with a mental imagery ROI because this area should theoretically (1) be more active during the imager’s than the aphantasic’s recall, given that the aphantasic has impaired mental imagery and (2) share some similarity between perception and recall, given that imagery is thought to involve some reactivation of perception. These were the same key hypotheses tested with the SVM searchlight. Therefore, to see whether we found the same trend of results, we averaged across the decoding accuracies for each voxel within the mental imagery ROI for each SVM searchlight condition. We only included voxels that were in the mask and inside the participants’ brains. Results for this analysis are reported in *Supplemental Results 2* and *Fig. S2*.

## Results

### Lower subjective imagery strength for aphantasic twin

The strength of mental imagery is often measured through self-report surveys, such as the VVIQ, which assesses the overall strength of mental imagery, and the Object-Spatial Inventory Questionnaire (OSIQ), which probes object and spatial imagery abilities separately. The imager twin reported scores within the standard imagery range (VVIQ=47, Object-OSIQ=56, Spatial-OSIQ=38). However, the aphantasic twin had scores indicative of overall diminished imagery and object imagery levels (VVIQ=24, Object-OSIQ=22), but intact spatial imagery (Spatial-OSIQ=49), as is typical for aphantasic individuals (Bainbridge, Pounder, et al., 2021).

Drawings made from memory have been shown to be a more objective measure of imagery experience (Bainbridge, Pounder, et al., 2021). When the twins drew three scene images from memory and perception (see *Drawing Experiment*), we observed the same trends as the reported survey results. Whereas both twins were able to accurately draw the scenes in detail during perception, the aphantasic twin used starkly less detail—including no color—when drawing the scenes from memory (**Fig. 2a**), despite the twins taking a comparable amount of time to produce these drawings from memory (aphantasic twin: *M* = 1 min 35 sec, imager twin: *M* = 2 min 3 sec). Both twins spent more time when drawing the images from perception (aphantasic twin: *M* = 7 min 25 sec, imager twin: *M* = 5 min 9 sec). Although the imager twin’s increased art experience (see *Table S1*) could influence these results, a large-scale study utilizing the same task still found that aphantasics drew significantly less from memory than imagers despite having the same level of artistic ability (Bainbridge, Pounder, et al., 2021). Therefore, lacking mental imagery likely reduces drawing performance during memory beyond what could be explained by artistic ability.

**Figure 2.**
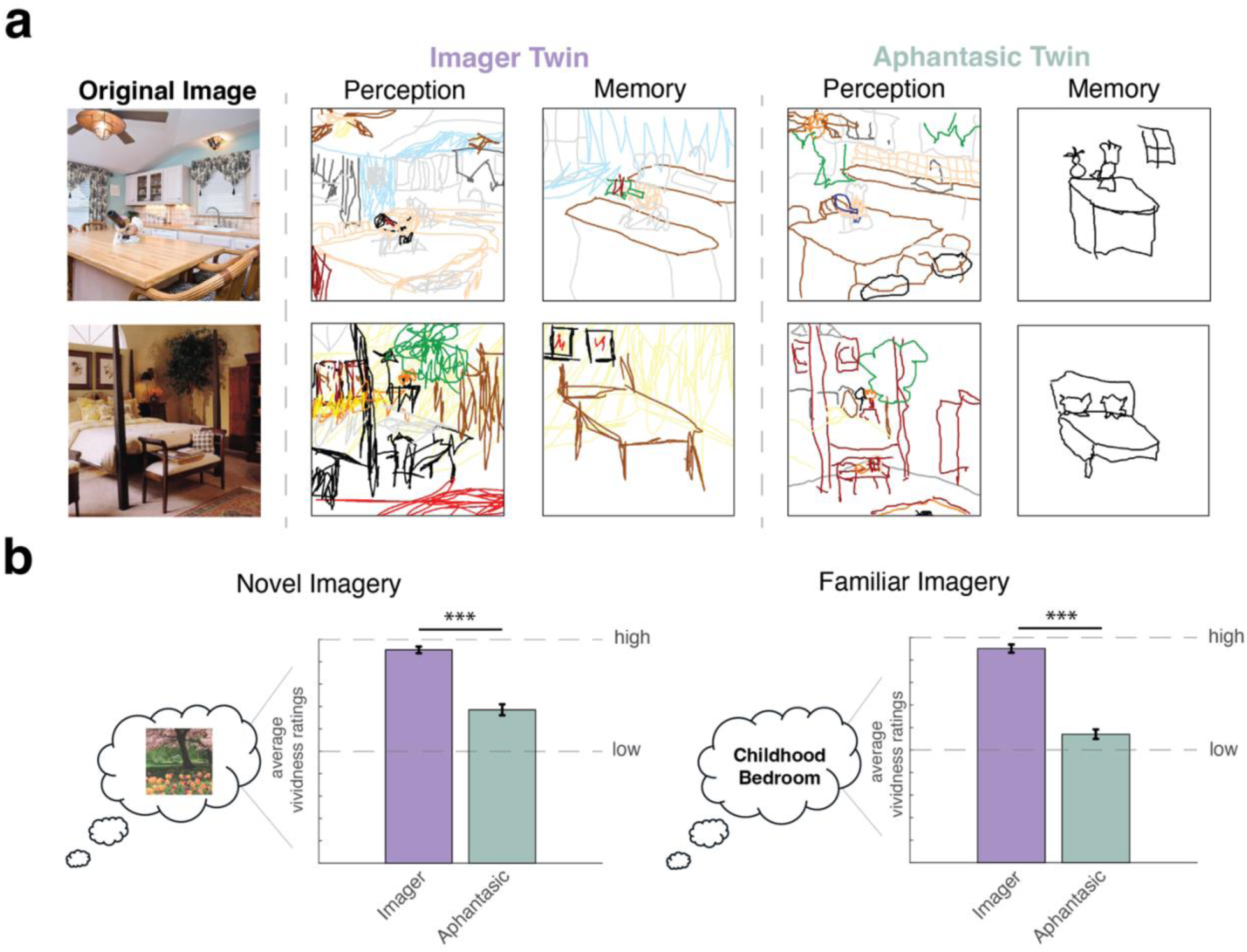
Behavioral results. (a) Drawings produced from memory and perception. Whereas both twins drew many objects in detail from a scene during perception, the aphantasic twin drew starkly less from memory compared to the imager twin. (b) Vividness responses during the imagery tasks. The aphantasic twin additionally reported significantly lower vividness during mental imagery for both the novel imagery and familiar imagery tasks.

Additionally, we analyzed the vividness reports collected after each in-scanner imagery trial (**Fig. 2b**). A 2-way ANOVA of successfully remembered trials with participant (imager/aphantasic) and task (novel imagery/familiar imagery) as factors revealed a significant effect of both participant (F(1,330)=246.68, p<0.001) and task (F(1,330)=8.44, p<0.004), as well as a significant interaction (F(1,330)=7.77, p=0.006). For novel images, the imager twin reported significantly higher imagery vividness for novel scenes and objects (*M*=1.91, *SD*=0.29) than the aphantasic twin (*M*=1.38, *SD*=0.49; t(190)=9.16, p<0.001). The imager twin also reported significantly higher vividness in her imagery for familiar people and places (*M*=1.90, *SD*=0.30) than the aphantasic twin *(M*=1.14, *SD*=0.35; t(140)=13.89, p<0.001). However, although the imager reported high vividness for both tasks—with no significant difference between tasks (t(148.91)=0.10, p=0.92)—the aphantasic twin actually reported significantly *lower* vividness during the familiar imagery than the novel imagery task (t(164.90)=3.61, p=0.003). This is opposite to the trend typically found in control participants (Ragni et al., 2021), suggesting that despite the twins having increased perceptual experience with familiar people and places than novel images, this experience does not benefit imagery in aphantasics like it does those with normal imagery.

### Similar univariate activation during imagery

Given the diminished mental imagery reported by the aphantasic twin, is there any information contained in her mental images? If there is category information during aphantasic imagery, then we should see differential univariate activation during imagery for different categories of items during the Novel Imagery task. Indeed, we observed higher activation for scenes than objects in scene-selective perceptual areas, such as the parahippocampal place area (PPA; **Fig. 3a**).

**Figure 3.**
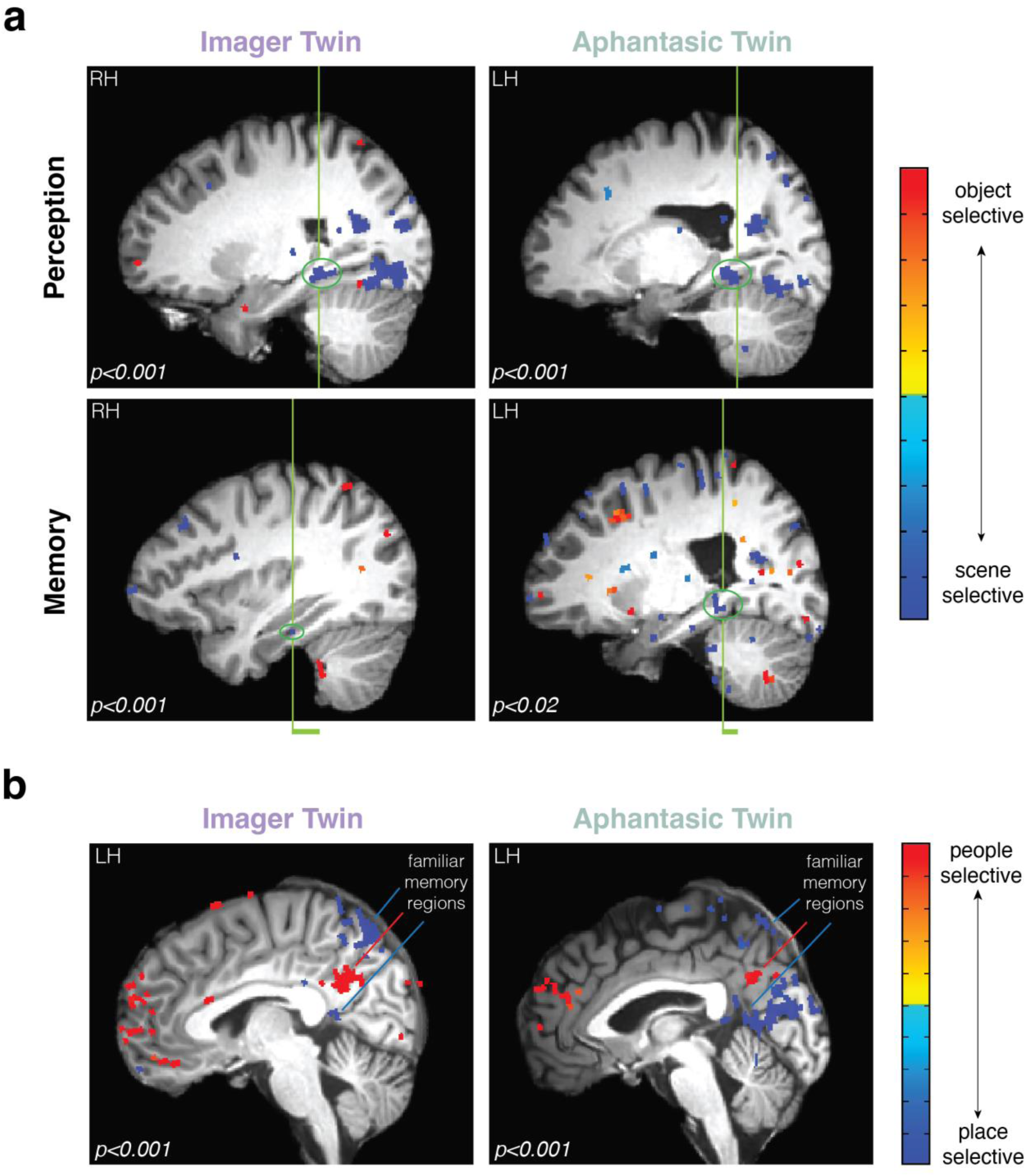
Univariate brain activity during the imagery tasks for both twins. (a) The location of PPA during perception and memory of the Novel Imagery task. The vertical green line indicates the location of the peak voxel activity in each condition. We observed an anterior shift in the peak voxel activity of PPA between perception and memory in both twins, with an equal (or even smaller) shift in the aphantasic compared to the imager. (b) A people>places contrast during the Familiar Imagery task. Using this contrast, we identified the recently discovered “familiar memory regions” in the medial parietal cortex in both twins, with their characteristic alternating pattern between familiar people and place selectivity. Each image is shown at a threshold of *p<0.001* unless otherwise noted, and all images are from the sagittal view. See also *Fig. S3* and *Table S3*.

We additionally looked at the location of these areas, with a focus on the PPA (**Fig. 3a**), as an anterior shift in peak voxel activity from perception to imagery (or memory) is thought to reflect the more conceptual nature of mnemonic compared to perceptual representations (Favila et al., 2020; Srokova et al., 2022). The peak voxel within the imager twin’s right PPA was anteriorly shifted from perception (*y*=29) to memory (*y*=34), although not for her left PPA (perception: *y*=30; imagery: *y*=30). However, we found a similar magnitude—or even smaller— of an anterior shift in the aphantasic’s left PPA between perception (*y*=26) and memory (*y*=28), suggesting items do not get more semanticized in aphantasic memory. We also found the anterior shift to be unilateral in the aphantasic twin, with no evidence of an anterior shift in her right PPA (perception: *y*=29; imagery: *y*=29). Interestingly, we did find very minimal evidence for a greater anterior shift in the aphantasic twin’s FFAs than the imager’s when comparing the peak voxel activity across tasks (see *Supplemental Results 3* and *Fig*. S4).

Are regions sensitive to the recall of familiar concepts also active during memory? Areas in the medial parietal cortex have been identified that alternate in their selectivity for familiar people and familiar places during imagery (Silson et al., 2019). To see if we could identify these areas in the aphantasic twin, we tested a univariate contrast of people>places in each twin (**Fig. 3b**). We found the characteristic alternating pattern of these familiar people and place memory regions in both twins, suggesting that information specific to familiar people and places are also present during aphantasic imagery. The locations of these areas aligned with where they have been found previously (Bainbridge & Baker, 2022; Silson et al., 2019).

### Different, though similar, brain patterns during imagery

As univariate activity revealed similarities—rather than differences—between the twins, what is causing their phenomenological differences? We hypothesized that these differences may be reflected in different *multivariate patterns* of activation during mental imagery, indicating different information stored in their mental images. To test this, we ran a whole-brain support vector machine (SVM) searchlight analysis on the neuroimaging data from the Novel Imagery task (**Fig. 4a**). To determine the similarity in representations across participants and across task phases, we trained an SVM to decode between objects and scenes in one condition (e.g., imager perception), and tested the decoding accuracy between objects and scenes in the other condition (e.g., aphantasic perception) within each searchlight region. To determine whether decoding was above chance, we ran a permutation test within each searchlight region.

**Figure 4.**
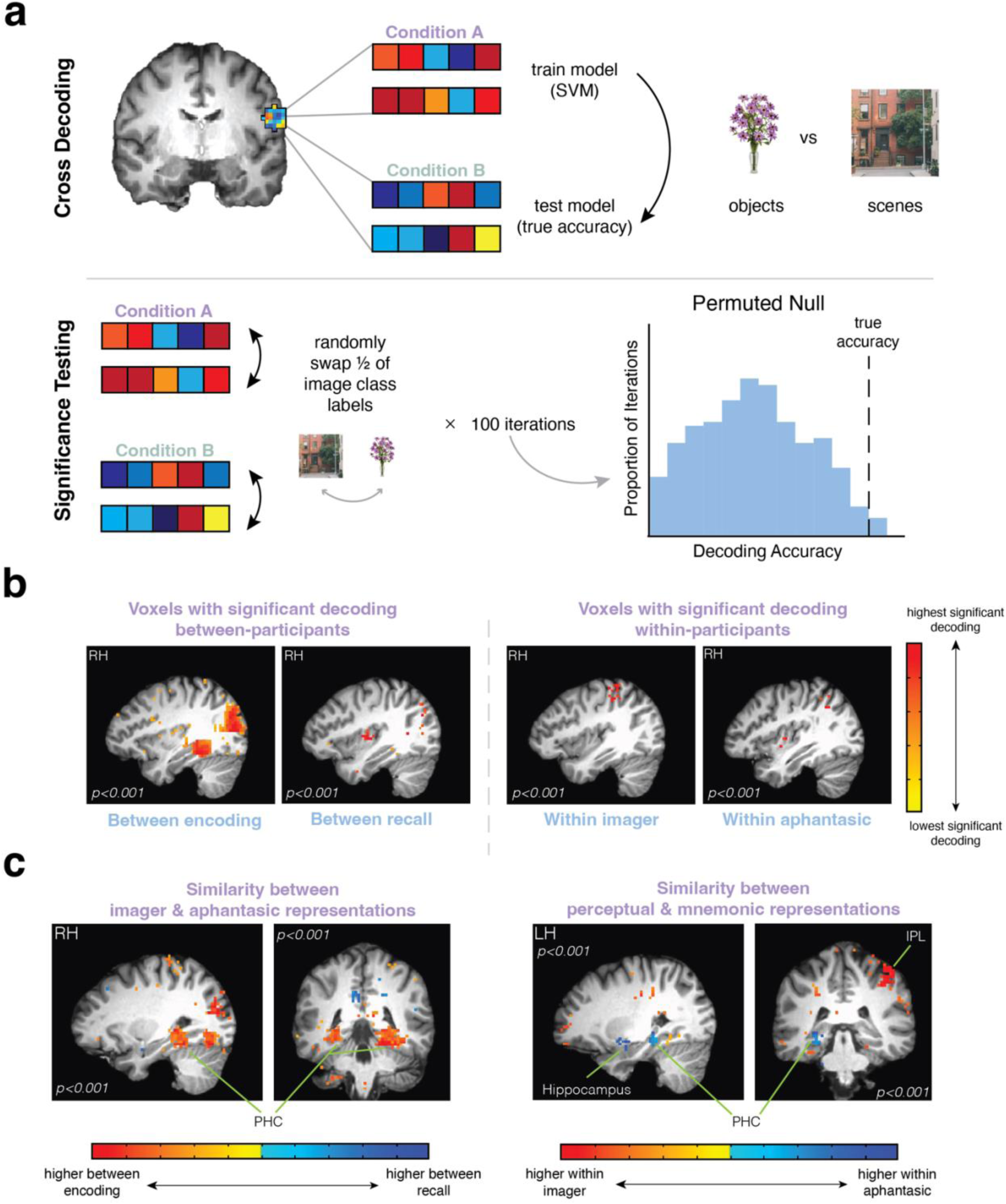
SVM searchlight methods and results. (a) Methods for cross decoding between conditions. Using the brain patterns within each searchlight region, we trained an SVM to distinguish between objects and scenes in one condition and tested on the other condition. Conditions were either between-participants (e.g., training on imager perception, testing on aphantasic perception) or within-participants (e.g., training on imager perception, testing on imager recall). To determine whether the voxels within a searchlight region were able to cross-decode above chance, we randomly swapped the image class labels for half of the training and test trials. We did this 100 times to build a null distribution to compare to the true decoding accuracy. (b) Voxels with significant decoding accuracy. Between-participants, there were many significant voxels able to cross-decode between the twins’ representations during perception, whereas there were far fewer during their recall. The decoding accuracy between the twins’ perceptual representations was also significantly higher than between their recall representations. Within-participants, there was a significantly higher decoding accuracy within the imager twin. However, the aphantasic twin had a surprisingly similar number of voxels as well as decoding accuracy as the imager twin. (c) Voxels with significantly higher decoding accuracy in one condition versus another. Whereas visual areas, including PHC, were significantly more similar between the twins’ perception than their recall, few areas emerged with higher similarity between their recall. Surprisingly, visual areas, including the PHC, shared significantly more similarity in their perceptual and mnemonic representations for the aphantasic than the imager. Each image is shown at a threshold of *p<0.001*, but all key regions reported survive cluster threshold correction (see *Supplemental Results 1* and *Fig. S1*). See also *Supplemental Results 2* and *Fig. S2* for an ROI-based approach.

First, given that aphantasics are thought to have intact perception but disrupted imagery, we hypothesized that there would be similarity between the twins’ perceptual representations, but not their recall representations (**Fig. 4b**). When we trained on the imager’s perceptual representations and tested on the aphantasic’s, we found a large number of voxels that were able to decode above chance (4511 voxels), with an average above-chance decoding accuracy of 66.8% (*SD*=5.5%). We also surprisingly found decodability between the twins’ recall representations, suggesting at least some shared information during imagery. However, this average decoding accuracy (*M*=60.7%, *SD*=2.0%) was significantly lower than between their perceptual representations (t(1192.95)=-48.64, p<0.001), and far fewer voxels were able to decode above chance (423 voxels).

Further, we hypothesized that if there is less perceptual information in aphantasic imagery, then there should be higher similarity between the imager twin’s perceptual and mnemonic representations than between the aphantasic’s (**Fig. 4b**). Within the imager, we found similarity between her perceptual and mnemonic representations, with 311 voxels able to decode above chance with 68.78% (*SD*=2.24) accuracy. However, we found a *surprisingly similar* degree of successful decoding between the aphantasic’s perceptual and mnemonic representations, with a similar number of significant voxels (263 voxels) and average decoding accuracy (67.89%, *SD*=2.50%). Although the imager twin’s decoding accuracy was significantly higher than the aphantasic’s (t(530.57)=4.49, p<0.001), the numerical difference of only ∼1% suggests that there might be more visual information present in memory for the aphantasic twin than we originally predicted. We replicated similar cross-decodability of perception and memory in both twins when targeting mental imagery areas as a region of interest (see *Supplemental Results 2* and *Fig. S2*).

What is the content of the information shared between conditions? To answer this, we first determined areas that had significantly higher decoding accuracy between the twins’ perceptual than between their recall representations using permutation testing (**Fig. 4c**). Many visual areas, including those extending along the parahippocampal cortex (PHC), had significantly higher decoding accuracy between the twins’ perceptual than between their recall representations. However, only a few areas—and none visual—had significantly higher decoding accuracy between their recall than between their perceptual representations. These results are in-line with what we would expect, with more visual information shared between the twins during perception than recall. Within-participants, given the lack of visual information in aphantasic memory, we predicted that visual memory areas would share significantly more information between perception and recall in the imager than the aphantasic twin. However, we surprisingly found evidence contrary to our prediction, with PHC and the hippocampus sharing significantly more similar representations during perception and memory in the aphantasic twin. As this posterior PHC region aligns with where the PPA is typically found (Epstein & Baker, 2019), this surprisingly suggests the presence of visual information in aphantasic memory. Given this intriguing finding, we visualized the imager’s and aphantasic’s representational structures during perception and memory of this task (see *Fig. S5*), but we did not find any conclusive evidence to explain these findings (i.e., no clear differences in representational patterns during memory). Within the imager, we found that the inferior parietal lobule (IPL) had significantly more similar representations during perception and memory in the imager than the aphantasic twin, which could suggest some immediate consolidation of visual information in the imager twin (Himmer et al., 2019).

### Different brain patterns during familiar imagery

Although we found evidence of visual information in memory for newly-learned images for the aphantasic twin, do we find this same evidence for more consolidated, highly familiar items? We tested this using the Familiar Imagery task—as this task required mentally imagining a personally familiar person or place *without* any perceptual information, the twins had to conjure visual detail from longer-term memory stores to accomplish this task. Indeed, the aphantasic twin’s lower vividness ratings compared to the Novel Imagery task supports the idea that this may be a more difficult task for those without imagery.

We constructed a representational similarity matrix (Kriegeskorte et al., 2008) by correlating brain activation between pairs of trials in the same PHC region that contained higher similarity between perception and memory in the aphantasic twin than the imager twin (**Fig. 5**). We quantified coarse level information by calculating discrimination indices (*D*), which subtracts the correlation *between* conditions (e.g., people and places) from the correlation *within* conditions (e.g., people and people). Therefore, if there is visual information in the aphantasic twin’s mental imagery when pulling from longer-term stores, then there should be positive discriminability for people versus places. However, we found a *D* close to 0 in the aphantasic twin, which was significantly lower than the positive *D* (0.098) in the imager twin (permutation testing: p<0.001). In other words, although this region contained category-level visual information for newly-learned images for the aphantasic twin, this visual information seemed to dissipate when drawing from longer-term memory stores.

**Figure 5.**
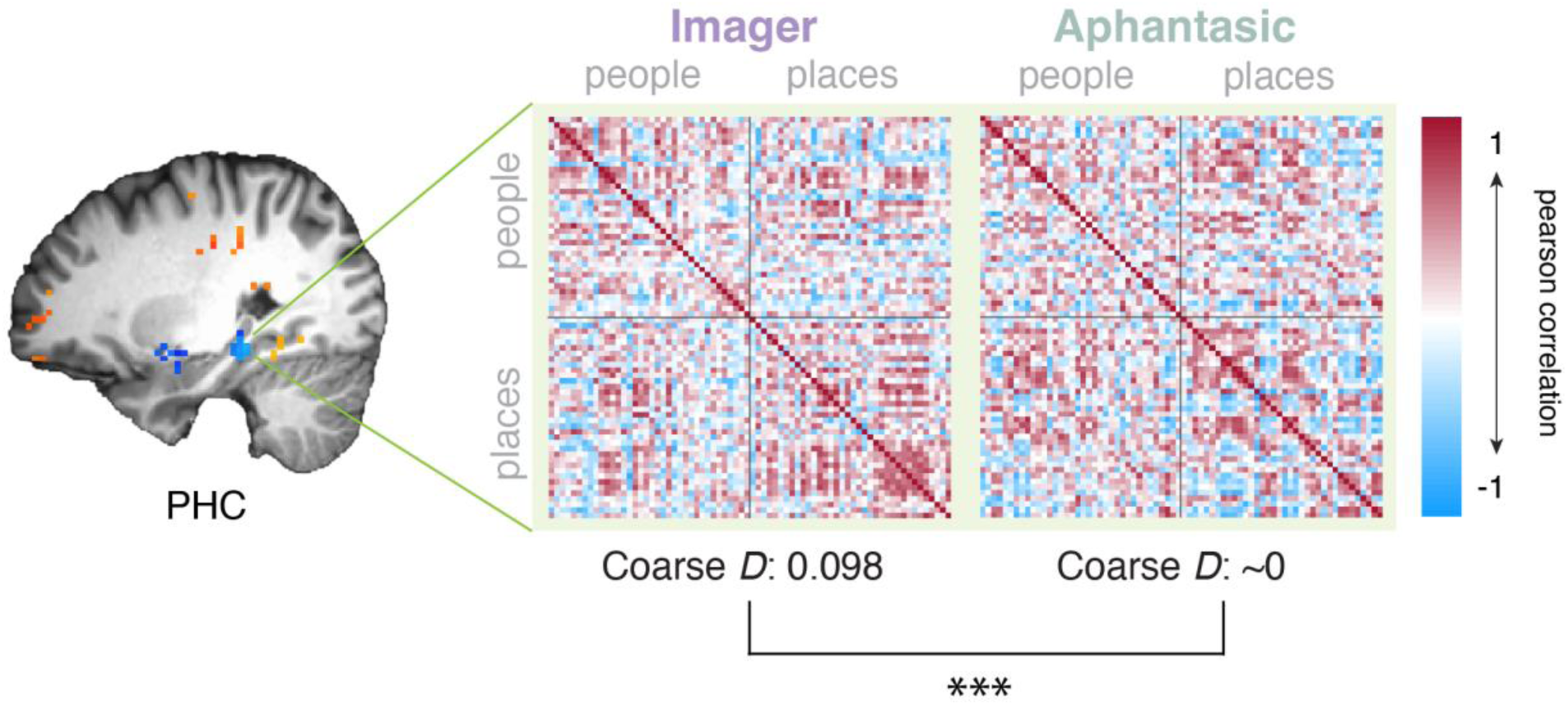
Representational similarity during familiar imagery. To determine whether there is coarse level (person vs. place) visual information in aphantasic memory during familiar imagery, we correlated brain activity from the PHC region between every pair of stimuli. We quantified the amount of coarse level information by calculating a discrimination index (D) for each twin, which subtracts the degree of neural similarity within category – between category. Although we found evidence of coarse level visual information in the imager twin, we found nearly next to no discrimination between people and places in the aphantasic twin. Indeed, D was significantly higher in the imager than the aphantasic twin. Pearson correlation values are shown here for visualization purposes, but all analyses were performed after these correlation values were Fisher Z-transformed.

### Lower and less visually-based connectivity between key regions in aphantasic twin

What could be the cause for the different amount of visual information in aphantasic memory between the two imagery tasks? To explore this, we quantified the strength of the twins’ functional connectivity during rest (**Fig. 6**). To compare the strength of their connections, we subtracted their correlation values between each pair of nodes (imager – aphantasic) and averaged across all the connections within lobes.

**Figure 6.**
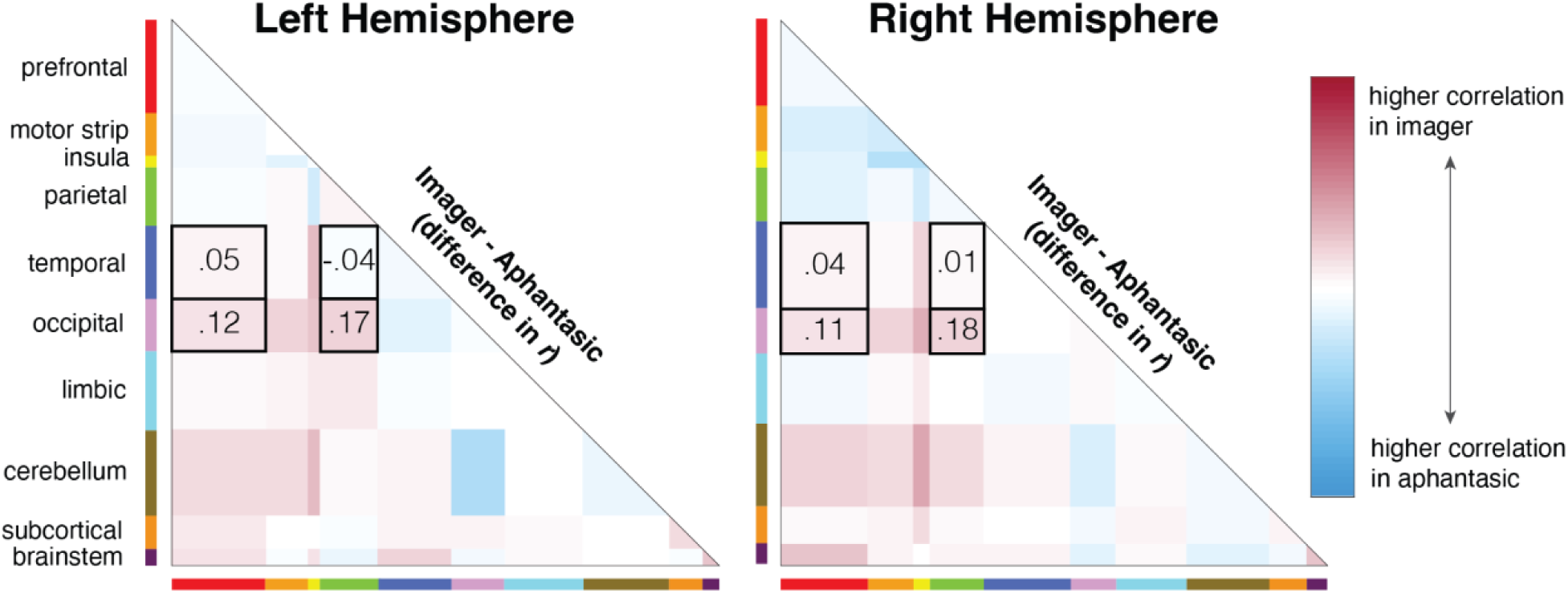
Differences in functional connectivity between the lobes of the brain. Red means a higher correlation between two lobes in the imager, whereas blue means higher correlation in the aphantasic. Interestingly, we generally found lower connectivity between lobes housing immediate memory processes (temporal and occipital) and lobes housing consolidated memory processes (parietal and prefrontal) in the aphantasic twin, which could account for the differences we found between imagery tasks. These connections of interest are outlined in black. See *Table S2* for all correlation values.

We overall found trends that replicate prior work, finding that the aphantasic twin had lower connectivity between her occipital lobe and both the prefrontal lobe (Liu et al., 2025; Milton et al., 2021) as well as the parietal lobe (Liu et al., 2025). However, we did not find lower connectivity between the occipital and temporal lobe as found previously (Zeman et al., 2010).

We additionally found that the aphantasic twin had reduced connectivity between her temporal lobe and both her prefrontal and parietal lobes (though only in the right hemisphere). Overall, the disconnect between occipitotemporal and fronto-parietal lobes in the aphantasic twin could interestingly hint at visual information initially making it into aphantasic memory, but unsuccessfully being consolidated into longer-term stores. All correlation values between lobes are reported in *Table S2*.

Additionally, we tested the *representational content* captured by these functional connections during rest, utilizing a model created by Ke et al. (in prep) that uses a participant’s functional connectivity patterns to determine the content of their mind-wandering. Ke et al. identified an “imagery network” whose strength predicts the degree to which an individual reports having thoughts in the form of images during rest. We calculated the mean strength of this network in each twin’s video watching data by subtracting functional connections that predicted less imagery from functional connections that predicted more imagery. The imagery network was significantly stronger in the imager twin (difference between networks*=*0.023) than the aphantasic twin (difference between networks=-0.025; *p*=0.010). These results suggest that the spontaneous thoughts of the aphantasic twin during rest may also lack visual imagery.

### Different language lateralization

Although identical twins raised in the same household are as similar as possible for two individuals, there is one notable physiological difference between these twins. Namely, whereas the imager twin self-reported as right-handed, the aphantasic twin self-reported as left-handed (called “mirror twins”). This opposite handedness was verified using the Edinburgh Handedness Inventory (EHI; imager twin EHI=1, aphantasic twin EHI=-1). As handedness can correlate with brain lateralization (Szaflarski et al., 2006), this meant that the laterality between the twins could also be opposite. We determined lateralization of the brain through a language localizer, in which the twins read words versus nonwords, to calculate a laterality index (LI). We found that the imager twin has left-side language localization (LI=0.225), but the aphantasic twin has bilateral dominance (LI=0.177), with a trend towards left-side language localization. As perceptual networks such as the face network have been found to be stronger in the right hemisphere (e.g., Lesinger et al., 2023), this means that the bilateral dominance of the aphantasic twin could distort and weaken the strength of these networks. Therefore, it is possible that the aphantasic twin’s different brain laterality could serve as a mechanism behind her lack of mental imagery.

## Discussion

In this work, we leveraged a rare sample of participants—identical twins, one with aphantasia and one with normal imagery—which allowed us to identify novel neural markers of aphantasia. First, we found similarity between the aphantasic and imager using univariate methods, with areas such as the posterior parahippocampal cortex (PHC) and “familiar memory regions” active during aphantasic memory. Second, when examining differences in multivariate patterns during memory between the twins, although we found significantly more similarity between the imager’s perceptual and mnemonic representations, we also found unexpected similarity between these representations for the aphantasic. In fact, visual areas in the PHC contained significantly higher similarity between the aphantasic’s perceptual and mnemonic representations than the imager’s. Although these findings suggest visual information in aphantasic memory, we did not find evidence for this during familiar imagery. Lastly, we found that the lack of visual information in aphantasic memory may be attributed to lower functional connectivity between occipitotemporal and frontoparietal areas.

As there have been fewer than a dozen published neuroimaging studies on aphantasia— and none looking at the *representational content* of aphantasic memory for higher-level features across the brain—the current results help build a foundation for our current understanding of the condition. In fact, even the discovery of these twins provides important insight into debates on whether aphantasia and imagery ability has a genetic component (Day et al., 2022; A. Zeman, 2024); since the twins have identical genetics, our study serves as evidence that aphantasia may not be fully determined by genetics. Further, our study provides evidence that there is indeed a difference between aphantasic memory content compared to controls, with our results suggesting that there is significantly less visual information in memory for both newly-learned images and familiar people and places. These neural results suggest that the lack of imagery is an objective experience, supporting other objective behavioral findings (Keogh & Pearson, 2018; Wicken et al., 2021), and that there may be ways to identify aphantasia on the neural level. The overall finding of less visual information in aphantasic memory also aligns with previous, more subjective measures of aphantasic memory content, such as recalling fewer objects and using less color when drawing scenes from memory (Bainbridge, Pounder, et al., 2021).

However, we also found evidence that memory content for the aphantasic may contain more visual information than we originally predicted. Although the difference in decoding accuracy between perception and memory in the imager twin was significantly higher than in the aphantasic twin, the accuracy was unexpectedly similar (only ∼1% difference). In fact, visual areas in the PHC had significantly *higher* decoding between perception and memory in the aphantasic twin, suggesting that there is still a surprising degree of perceptual information in aphantasic memory for newly-learned images. Univariate approaches also revealed intact category-level visual information for the aphantasic, with activation of PPA, OPA, and LO during memory for newly-learned images. We even found activation of regions selective to recall of familiar people and places (“familiar memory regions”), suggesting at least some intact memory content during familiar imagery as well.

The finding of less visual information in aphantasic memory during familiar imagery compared to novel imagery also suggests that the amount of visual information may depend on when the information was learned. Whereas novel imagery involved mentally imagining an image that was shown shortly before, familiar imagery required mentally imagining a familiar person or scene without any previous visual information shown. Therefore, it is possible that aphantasics can maintain some visual information shortly after encoding, but that this visual information dissipates more in aphantasics than imagers when the memory becomes consolidated. Indeed, upcoming work may support this idea, which reports aphantasics maintaining visual information in early visual cortex during working memory (Weber et al., 2024).

Additionally, our results may suggest that there is a transformation between perception and memory representations, even for those with normal imagery. As the aphantasic twin’s memory lacks perceptual information, it likely undergoes a transformation in representation from perception (e.g., becomes more semanticized). As a result, their memory serves as a powerful comparison to determine whether such a transformation occurs even for those who report more visual information in memory. Since we found that there was comparable cross-decoding between perception and memory between the twins, this suggests the removal of some perceptual information—and thereby a transformation—in the imager’s memory as well. Similarly, we also found a similar degree of an anterior shift in peak voxel activity for areas like PPA between the imager and aphantasic. These results therefore align with previous studies that have found differences between perception and memory (Bainbridge, Hall, et al., 2021; Baldassano et al., 2016; Silson et al., 2019; Steel et al., 2021).

Likewise, our results also support theories of mental imagery that go beyond pure perceptual reinstatement. For example, “vision in reverse” proposes that voluntary mental imagery constructs an image by combining stored memories—meaning that we can form novel images never perceived in the real world—with this process beginning in the frontal cortex before moving to the medial temporal lobe and ‘backwards’ to the visual cortex (Dentico et al., 2014; Dijkstra et al., 2017; Pearson, 2019). Given that we found reduced connectivity between the aphantasic twin’s prefrontal and temporal lobes, it is therefore plausible that aphantasics may lack the ability to initiate this “vision in reverse” process, leading to diminished mental imagery. Further, we found no evidence for retinotopic reinstatement (Albers et al., 2013; Cichy et al., 2012; Lee et al., 2012), with no cross-decoding between perception and imagery in low-level visual areas like primary visual cortex. However, it is possible that this is due to our imagery tasks involving mentally imagining whole images of places, people, and objects rather than stimuli focusing more on low-level features, like gratings (Kosslyn & Thompson, 2003).

Our network results also provide an important replication of previous network-based studies. Specifically, we found both lower connectivity between the aphantasic twin’s occipital lobe and her prefrontal (Liu et al., 2025; Milton et al., 2021) and parietal lobes (Liu et al., 2025). However, we did not find lower connectivity between the aphantasic’s occipital and temporal lobe as reported previously (Zeman et al., 2010), and we found a new pattern of lower connectivity between the aphantasic’s temporal lobe with her prefrontal and parietal lobes. In total, these trends in connectivity in the aphantasic suggest that there could be a lack of access to visual information for consolidated memories. Indeed, the occipital lobe is widely known to house visual areas like OPA (Dilks et al., 2013) and the temporal lobe to house the hippocampus and other visual areas like PPA (Epstein & Kanwisher, 1998). In contrast, the medial prefrontal cortex and posterior parietal cortex are thought to be two areas that house longer-term stores in the greater neocortex after memory consolidation (Sekeres et al., 2018; Tompary & Davachi, 2017), which involves migration of memories from the hippocampus (Himmer et al., 2019). Therefore, it is possible that more visual information is lost during consolidation for aphantasics than imagers.

Lastly, the present study raises important avenues of research for future work. Although the twins are identical, we did find that the imager twin processed language in the left hemisphere, whereas the aphantasic twin processed language bilaterality (though left-hemisphere leaning). As visual processing is more right lateralized (Hugdahl, 2011; Hugdahl & Davidson, 2003), the aphantasic twin having a less typically lateralized brain could mean that some of her visual processing is inhibited. Therefore, future work could investigate whether there is a connection between brain laterality and imagery ability. In addition, the presence of visual information in aphantasic memory in the present study suggests that aphantasia may be more of a spectrum than a discrete condition. Indeed, although the aphantasic twin reported imagery within the aphantasic range, they did report some visual information in imagery. Therefore, it may be valuable for future studies investigating aphantasia to recruit participants with the lowest VVIQ score, indicating the complete absence of visual imagery.

In conclusion, this case study of identical twins was able to characterize aphantasia in new and valuable ways, quantifying their memories as lacking visual information even on the most objective, neural level. However, this study also revealed that there can still be a surprising level of visual information for someone with aphantasia, at least for newly-learned images.

## Acknowledgements

We are immensely grateful to Jin Ke and Monica Rosenberg for allowing us to use their method to quantify the content of the twins’ mind-wandering to include in this paper. We would additionally like to thank Hayoung Song and Ziwei Zhang for their help with the functional connectivity analysis. The authors would like to thank the MRI Research Center, the University of Chicago (MRIRC, RRID:SCR_024723) for their assistance in MRI data acquisition of this study. This work was supported by the National Eye Institute (R01-EY034432) to W.A.B.

