## Supplementary Materials for "The Neural Underpinnings of Aphantasia: A Case Study of Identical Twins"

* These authors contributed equally

Correspondence to: Emma Megla

940 E. 57th St.

Chicago, IL 60637

<u>Contents</u> <u>Page</u>

**Supplemental Table 1:** Interview questions and answers assessing the natural history of the twins 3

**Supplemental Results 1:** Different, though similar, brain patterns during imagery using cluster threshold correction 4

**Supplemental Figure 1:** SVM searchlight results with cluster threshold correction 5

**Supplemental Results 2:** SVM cross-decoding in mental imagery region of interest 6

**Supplemental Figure 2:** Decoding accuracy in mental imagery region 7

**Supplemental Table 1:** Differences in correlation values (imager – aphantasic) across all lobes of the brain 8

**Supplemental Table 2:** MNI coordinates for category-selective areas during perception and memory 9

**Supplemental Figure 3:** Peak voxel activity in the medial temporal lobe (MTL) during perception and memory for the aphantasic twin 10

**Supplemental Results 3:** Peak voxel activity in fusiform face area (FFA) during perception and familiar imagery 11

**Supplemental Figure 4:** The location of FFA during perception and familiar imagery 12

**Supplemental Figure 5:** Representational similarity matrices (RSMs) during encoding and recall in the novel imagery task 13

**Table S1. Interview questions and answers assessing the natural history of the twins**

| Question | Abbreviated Imager Response | Abbreviated Aphantasic Response |
|---|---|---|
| **Describe your imagery experience.** | visual person; communicates imagery with clients | intact imagery in other senses besides vision |
| **When did you become aware of the difference in your imagery abilities?** | college | was 29 or 30 when came across aphantasia article |
| **What is your occupation?** | graphic designer | chemist; chemistry and engineering research technician |
| **If you're willing to answer, what has your health history been like?** | ovarian cancer (6 months); anxiety | lead poisoning as kids (both twins); anxiety; depression; ADHD |
| **What is your relationship status?** | married (2 years); in relationship (10 years) | married (6 years) |
| **What type of books, tv, and movies do you like (if any)?** | animated TV and movies; fiction podcasts; visually stimulating pieces of media; strategy and role-playing games | sci-fi and fantasy books and video games; mystery TV |
| **What are your hobbies?** | calligraphy; journaling; drawing; traveling; getting manicures | rock climbing; reading; knitting; gardening; video games |
| **Do you have any prior art experience?** | fine arts bachelor's degree | painted as a kid; no formal training |
| **Do you have any strategies to compensate for not being able to form clear mental images?** | N/A | writing; other senses |
| **Have you noticed any differences in your language abilities or any other cognitive abilities?** | aphantasic twin needs to sketch things out; aphantasic twin has hard time planning routes when rock climbing | aphantasic twin worse memory |

**Supplemental Results 1. Different, though similar, brain patterns during imagery using cluster threshold correction**

As the data and results in the primary paper are reported at an uncorrected threshold of *$p<0.001$*, here we present the SVM searchlight results after cluster threshold correction (see *Whole-brain SVM searchlight analyses*), which mirror the same trends and the same key regions emerge. First, when we trained on the imager's perceptual representations and tested on the aphantasic's, we found many voxels able to decode above chance (4179 voxels) with a high decoding accuracy ($M$=67.33%, $SD$=5.37%; see **Fig. S1a**). However, we found fewer voxels able to decode between the twins' recall representations (55 voxels), and with significantly lower decoding accuracy ($M$=62.22%, $SD$=1.75%; $t(1192.90)=48.64$, $p<0.001$). Next, when we trained on perception and tested on recall within the same twin, we found 104 voxels able to decode above chance in the imager with an average decoding accuracy of 69.55% ($SD$=1.84%). There were slightly fewer voxels able to decode above chance in the aphantasic (75 voxels), and there was significantly lower decodability between perception and memory in the aphantasic twin ($M$=69.06%, $SD$=2.45%; $t(530.57)=4.49$, $p<0.001$). However, same as the results reported in the primary paper, the aphantasic still had surprisingly high decoding accuracy between perception and memory, with a numerical difference of less than 1% between the twins.

When we isolated areas that had significantly higher decoding accuracy in one SVM condition than the other (either between- or within-participants), we found the same key regions emerge as the uncorrected results in the primary paper (see **Fig. S1b**). When comparing perception to memory, we found higher similarity in visual areas between the twins' perceptual representations than between their mnemonic. However, there were few areas (and none visual) that contained higher similarity between the twins' mnemonic representations than their perceptual representations. When comparing areas that had significantly higher decoding accuracy between perception and memory in one twin, we found that the inferior parietal lobule (IPL) contained more similar representations in the imager. Surprisingly, we found that the hippocampus and parahippocampal cortex (PHC) contained more similarity between perception and memory in the aphantasic twin, suggesting at least some visual information in aphantasic memory.

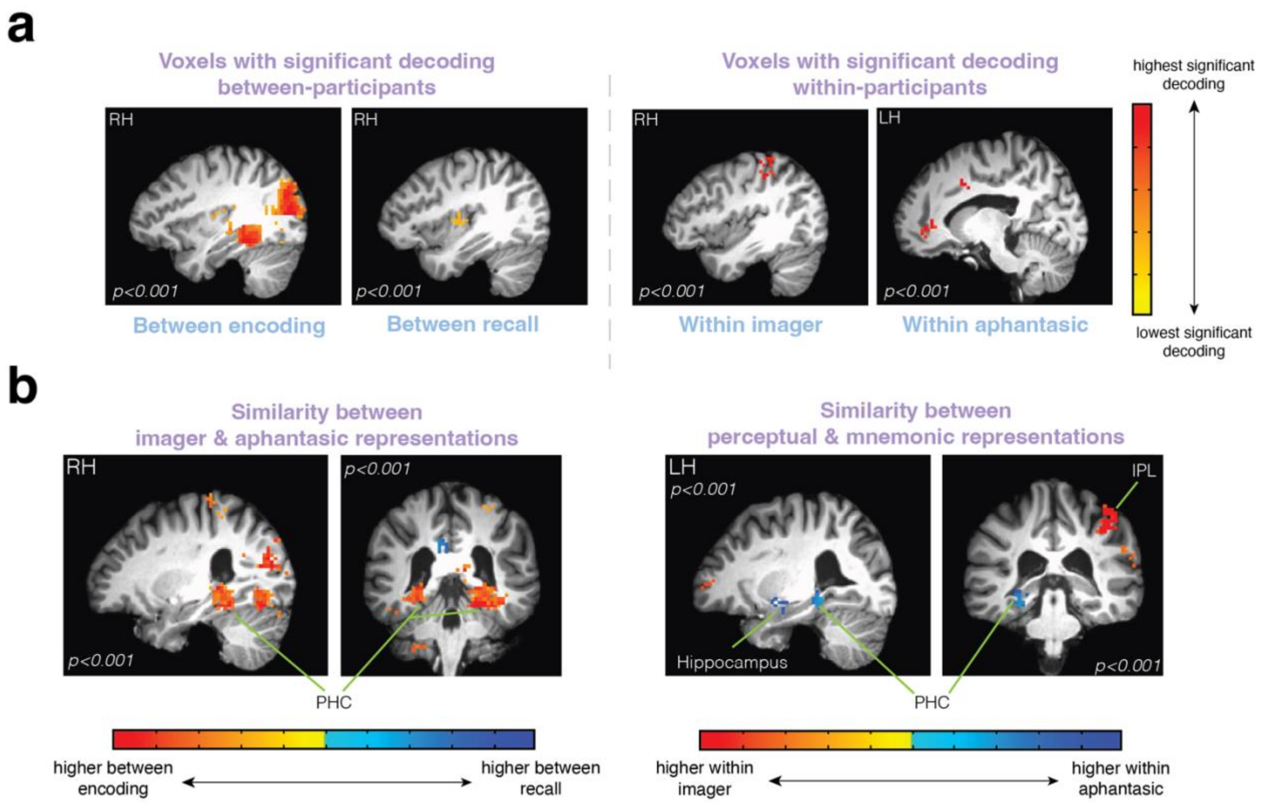


**Figure S1. SVM searchlight results with cluster threshold correction.** (a) Voxels with significant decoding accuracy after cluster threshold correction for each SVM searchlight condition. Between-participants, we found many more voxels able to decode between the twin's encoding than between their recall representations, and with significantly higher decoding accuracy. Within-participants, there were slightly more voxels able to decode between perception and memory within the imager, and with significantly higher decoding accuracy, than within the aphantasic twin. However, there was surprisingly high decodability within the aphantasic. (b) Voxels with significantly higher decoding accuracy in one condition than the other after cluster threshold correction. When comparing perception to memory, more visual areas, such as PHC, contained higher similarity between the twins' perceptual representations than between their mnemonic representations. Surprisingly, when comparing areas that contained significantly more similar representations between perception and memory in one twin, visual area PHC had significantly more similar representations in the aphantasic twin. All results and key regions identified replicate those found in the primary paper.

**Supplemental Results 2. SVM cross-decoding in mental imagery region of interest**

To determine whether the SVM searchlight results can be replicated in an ROI analysis, we calculated the average decoding accuracy for each SVM searchlight condition within a mental imagery ROI (see *Mental imagery ROI SVM analysis*). Overall, we found the same trends both between-participants and within-participants as the initial SVM searchlight analysis (see **Fig. S2**). When testing the similarity in representations between the twins in this mental imagery area, we found that there was significantly higher decoding accuracy between their perceptual representations (*M*=54.19, *SD*=8.44) than between their recall representations (*M*=49.28, *SD*=4.80; independent samples *t*-test: $t(2080)=16.31$, $p<0.001$). We also found that the imager twin had a significantly higher decoding accuracy between their perceptual and memory representations (*M*=55.77, *SD*=4.58) than the aphantasic twin (*M*=55.00, *SD*=4.80; independent samples *t*-test: $t(2080)=3.75$, $p<0.001$). In fact, similar to the SVM searchlight analysis, although this difference between the twins was significant, the decoding accuracy was comparable and less than 1% different. Therefore, both the SVM searchlight results and the current ROI analysis suggest that though there is significantly less visual information in aphantasic memory, there is more visual information than we originally anticipated.

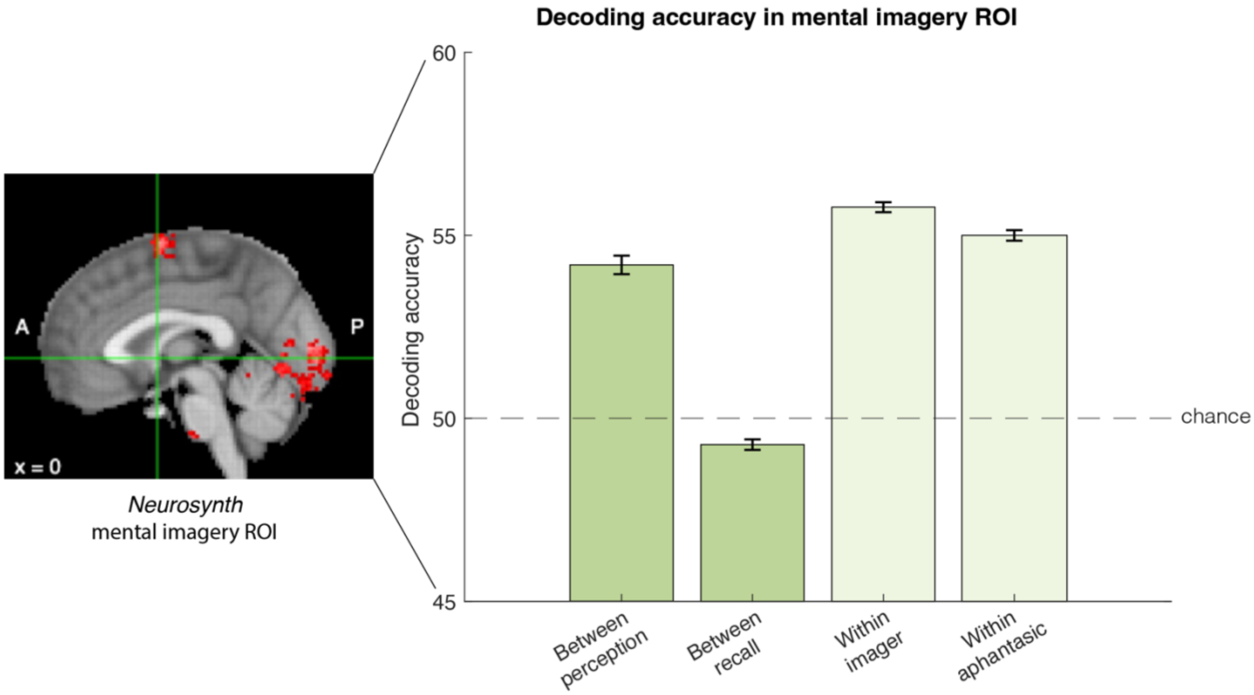


**Figure S2. Decoding accuracy in mental imagery regions.** A "mental imagery" region of interest (ROI) was isolated using *Neurosynth*, which compiled voxels that have preferentially been found to be active during mental imagery across 84 studies. For each SVM condition, we averaged across the decoding accuracies of every voxel in this mental imagery ROI. The overall trends of these results replicate the SVM searchlight analyses both between- and within-participants. Between-participants, there was a numerically higher similarity between the twins' perceptual than recall representations. Within-participants, there was a numerically higher decoding accuracy between perception and memory within the imager, though with surprisingly similar decoding accuracy within the aphantasic.

**Table S2. Differences in correlation values (imager – aphantasic) across all lobes of the brain**

| lobe | pre-frontal | motor strip | insula | parietal | temporal | occipital | limbic | cerebellum | sub-cortical | brain-stem |
|---|---|---|---|---|---|---|---|---|---|---|
| *Left Hemisphere* | | | | | | | | | | |
| prefrontal | -.04 | | | | | | | | | |
| motor strip | -.06 | -.03 | | | | | | | | |
| insula | -.06 | -.13 | -.05 | | | | | | | |
| parietal | -.05 | .01 | -.22 | .06 | | | | | | |
| temporal | .05 | .01 | .27 | -.04 | -.08 | | | | | |
| occipital | .12 | .18 | .22 | .17 | -.14 | -.06 | | | | |
| limbic | .01 | .04 | .07 | .09 | -.05 | -.01 | -.02 | | | |
| cerebellum | .15 | .15 | .26 | .02 | .05 | -.35 | -.02 | -.11 | | |
| subcortical | .07 | -.02 | -.02 | -.06 | .05 | .06 | .03 | -.02 | .14 | |
| brainstem | .11 | -.06 | .11 | -.10 | .16 | -.09 | .01 | -.01 | -.04 | .24 |
| *Right Hemisphere* | | | | | | | | | | |
| prefrontal | -.08 | | | | | | | | | |
| motor strip | -.18 | -.20 | | | | | | | | |
| insula | -.14 | -.34 | -.33 | | | | | | | |
| parietal | -.14 | -.08 | -.20 | -.08 | | | | | | |
| temporal | .04 | .02 | .18 | .01 | -.03 | | | | | |
| occipital | .11 | .18 | .33 | .18 | -.03 | .02 | | | | |
| limbic | -.07 | .02 | .09 | -.02 | -.07 | .01 | -.04 | | | |
| cerebellum | .16 | .14 | .35 | .14 | .04 | -.18 | .01 | -.07 | | |
| subcortical | .01 | .04 | .13 | .02 | -.02 | -.08 | .06 | -.08 | .04 | |
| brainstem | .22 | .05 | <-.01 | .06 | .05 | -.13 | <.01 | -.15 | -.09 | .22 |

**Table S3. MNI coordinates for category-selective areas during perception and memory**

| | | *Imager* | | | |
|---|---|---|---|---|---|
| | | *left hemisphere* | | *right hemisphere* | |
| | | perception | memory | perception | memory |
| *Parahippocampal Place Area (PPA)* | (*x*,*y*,*z*) | (19,30,19) | (18,30,19) | (38,29,21) | (42,34,17) |
| | *p*-threshold | *p<0.001* | *p<0.001* | *p<0.001* | *p<0.001* |
| *Medial Place Area (MPA)* | (*x*,*y*,*z*) | (25,25,27) | (24,21,30) | (33,26,26) | (35,23,29) |
| | *p*-threshold | *p<0.001* | *p<0.001* | *p<0.001* | *p<0.001* |
| *Occipital Place Area (OPA)* | (*x*,*y*,*z*) | (18,14,35) | (17,17,32) | (40,13,32) | (44,15,34) |
| | *p*-threshold | *p<0.001* | *p<0.001* | *p<0.001* | *p<0.001* |
| *Lateral Occipital (LO)* | (*x*,*y*,*z*) | (15,20,21) | (13,20,22) | (47,19,19) | (46,56,69) |
| | *p*-threshold | *p<0.001* | *p<0.001* | *p<0.001* | *p<0.001* |

| | | *Aphantasic* | | | |
|---|---|---|---|---|---|
| | | *left hemisphere* | | *right hemisphere* | |
| | | perception | memory | perception | memory |
| *Parahippocampal Place Area (PPA)* | (*x*,*y*,*z*) | (24,26,21) | (23,28,21) | (39,29,22) | (42,29,20) |
| | *p*-threshold | *p<0.001* | *p<0.02* | *p<0.001* | *p<0.011* |
| *Medial Place Area (MPA)* | (*x*,*y*,*z*) | (27,26,26) | (24,22,31) | (34,24,29) | (36,23,31) |
| | *p*-threshold | *p<0.001* | *p<0.001* | *p<0.001* | *p<0.001* |
| *Occipital Place Area (OPA)* | (*x*,*y*,*z*) | (22,14,35) | (21,14,31) | (41,15,34) | (42,14,28) |
| | *p*-threshold | *p<0.001* | *p<0.001* | *p<0.001* | *p<0.001* |
| *Lateral Occipital (LO)* | (*x*,*y*,*z*) | (14,19,22) | (15,24,27) | (47,20,18) | (41,17,26) |
| | *p*-threshold | *p<0.001* | *p<0.001* | *p<0.001* | *p<0.001* |

*Note*. p-threshold is the lowest p-value the region was able to be identified

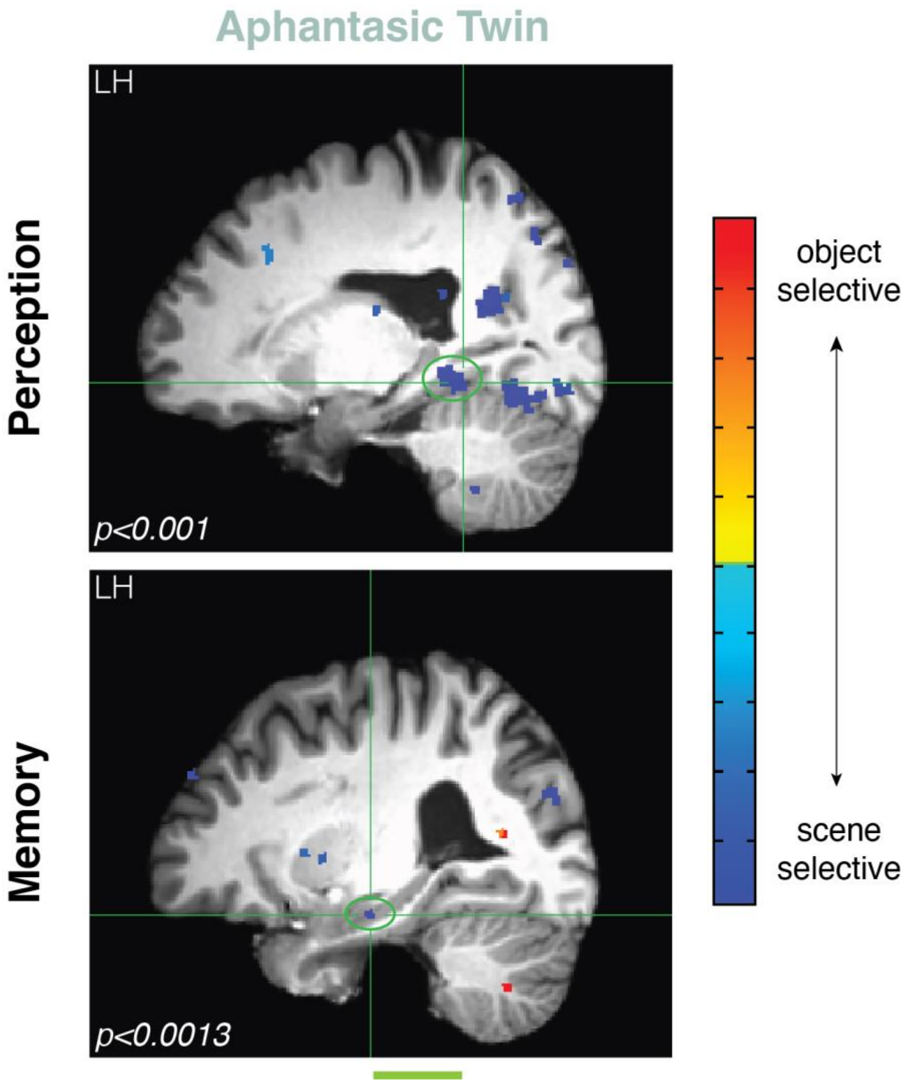


**Figure S3. Peak voxel activity in the medial temporal lobe (MTL) during perception and memory for the aphantasic twin.** Due to the liberal threshold needed to identify PPA during memory for the aphantasic twin, we also identified peak voxel activity in the greater MTL using a more conservative threshold. Using this criteria, we found a similar magnitude of an anterior shift (or larger) between perception (*y*=26) and memory (*y*=37) in the aphantasic twin's MTL than the imager twin's.

**Supplemental Results 3. Peak voxel activity in fusiform face area (FFA) during perception and familiar imagery**

Although the twins never perceived and mentally imagined the same faces during the experiment as they did for scenes and objects, they did perceive faces during the perceptual localizer and imagined familiar faces during the Familiar Imagery task. Therefore, we can use this separate data to determine whether there was an anterior shift in peak voxel activity in FFA—like we observed in PPA—in either twin. Note that although FFA is classically defined as a functional region sensitive to face perception, here we additionally use "FFA" to also refer to nearby regions that are sensitive to faces during memory and imagery. The FFA was localized using a person > place contrast in both tasks (see **Fig. S4**).

In the imager twin, we found no evidence of an anterior shift from face perception to face imagery in either the left FFA (perception: $x = 18$, $y = 23$, $z = 19$; memory: $x = 18$, $y = 22$, $x = 20$) or the right FFA (perception: $x = 44$, $y = 25$, $z = 18$; memory: $x = 46$, $y = 24$, $z = 17$). However, we did find evidence for a very minor anterior shift in both the left (perception: $x = 13$, $y = 23$, $z = 16$; memory: $x = 15$, $y = 24$, $x = 21$) and right FFA (perception: $x = 43$, $y = 22$, $z = 18$; memory: $x = 45$, $y = 23$, $x = 17$) in the aphantasic twin. Therefore, in contrast to many of the univariate analyses reported in the main text, these results could suggest minor univariate differences between the twins. Specifically, it is possible that the anterior shift observed in the aphantasic twin, but not the imager twin, could support our hypothesis of a greater semanticization in those with aphantasia. However, given that the observed anterior shift in both the left and right FFA were only one voxel, and that this analysis was conducted across different data (novel faces in perception versus familiar faces in memory), future work should investigate whether there is an anterior shift in FFA for aphantasics but not imagers using a larger sample size of participants.

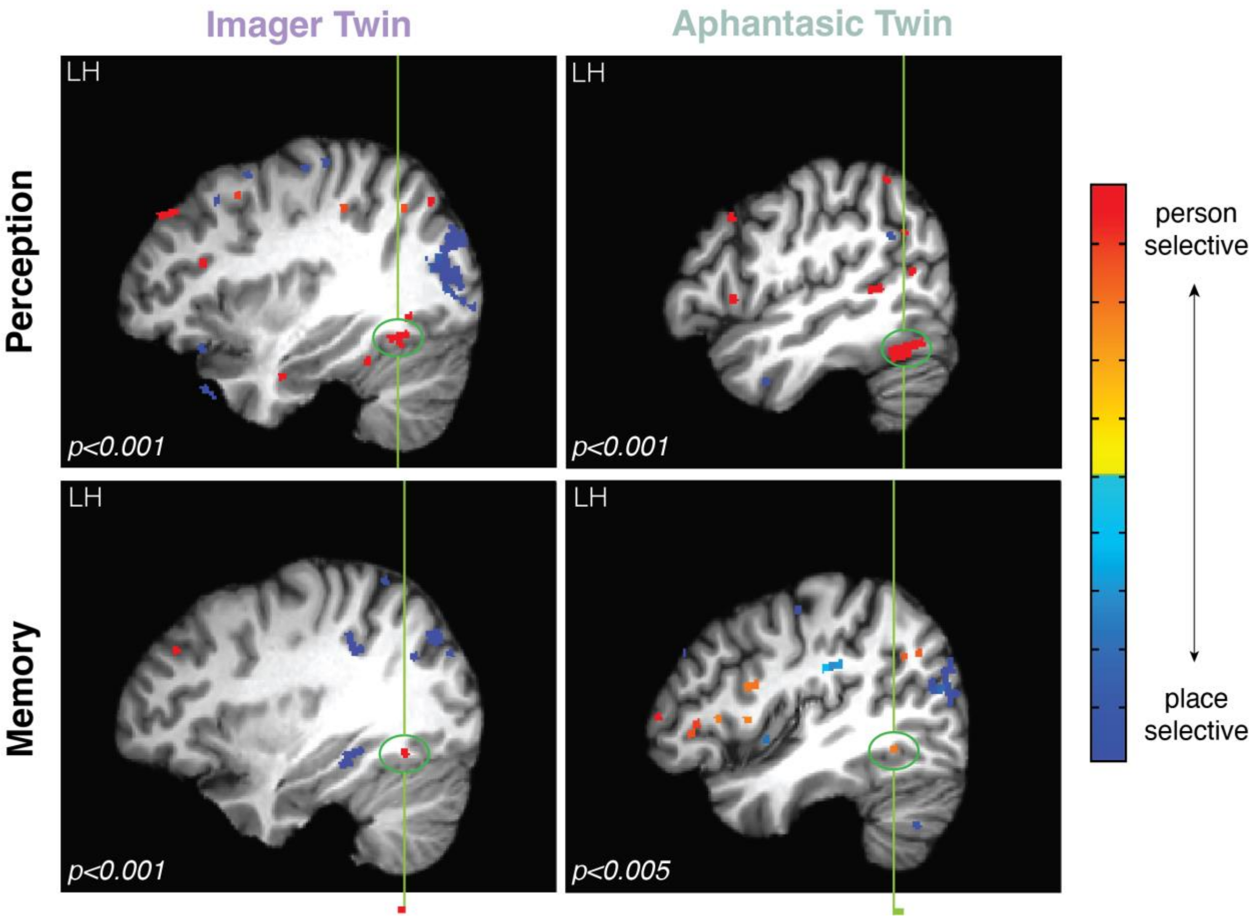


**Figure S4. The location of FFA during perception and familiar imagery.** Peak voxel activity was determined for perception using data from the perceptual localizer and was determined for memory using data from the Familiar Imagery task. The green vertical line indicates the location of this peak voxel activity in each condition. We found very minimal evidence for an anterior shift in the aphantasic twin, but not the imager twin. While only results from the left hemisphere are shown here, we found the same trends in the right hemispheres of each twin. Each image is shown at a threshold of $p<0.001$ unless otherwise noted, and all images are from the sagittal view.

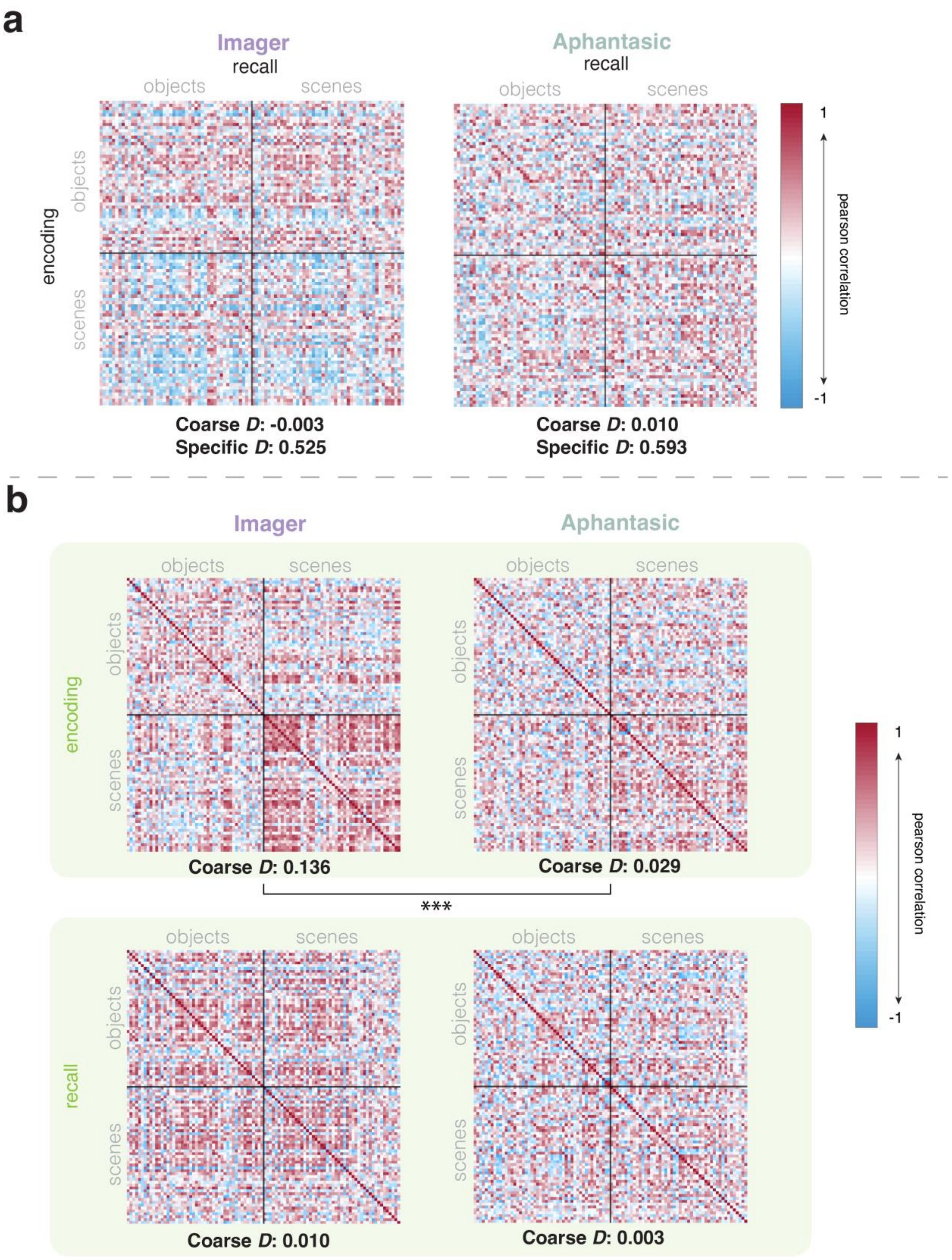
a
Imager
recall
objects
scenes
encoding
objects
scenes
Coarse D: -0.003
Specific D: 0.525
Aphantasic
recall
objects
scenes
Coarse D: 0.010
Specific D: 0.593
1
pearson correlation
-1
b
Imager
Aphantasic
encoding
objects
scenes
Coarse D: 0.136
Coarse D: 0.029
***
recall
objects
scenes
Coarse D: 0.010
Coarse D: 0.003
1
pearson correlation
-1

**Figure S5. Representational similarity matrices (RSMs) during encoding and recall in the novel imagery task.** (a) RSMs between encoding and recall within each twin. Coarse discrimination indices (*D*) were calculated as within-category similarity — between-category similarity on Fisher Z-transformed Pearson correlation values, and significance was determined through permutation testing of these values. Specific discrimination indices (*D*) were calculated as similarity along the diagonal — within-category similarity. Pearson correlation values are shown in the RSMs. Although we replicate the pattern found from the SVM analysis of higher coarse discriminability between encoding and recall in the aphantasic twin, this did not reach significance when comparing the Coarse discrimination indices of the twins. Interestingly, we found evidence for fine-grained information in memory with positive Specific discrimination indices for both twins. (b) Separate RSMs for encoding and recall. We found significantly higher discrimination between scenes and objects during encoding in the imager twin, potentially suggesting they had stronger memory traces during encoding. However, we did not find any significance difference in discriminability between the twins during recall.
